# Genomic dissection of bipolar disorder and schizophrenia including 28 subphenotypes

**DOI:** 10.1101/173435

**Authors:** Douglas M Ruderfer, Stephan Ripke, Andrew McQuillin, James Boocock, Eli A Stahl, Jennifer M Whitehead Pavlides, Niamh Mullins, Alexander W Charney, Anil P S Ori, Loes M Olde Loohuis, Enrico Domenici, Arianna Di Florio, Sergi Papiol, Janos L. Kalman, Rolf Adolfsson, Ingrid Agartz, Esben Agerbo, Huda Akil, Diego Albani, Margot Albus, Martin Alda, Madeline Alexander, Judith Allardyce, Ney Alliey-Rodriguez, Thomas D Als, Farooq Amin, Adebayo Anjorin, Maria J Arranz, Swapnil Awasthi, Silviu A Bacanu, Judith A Badner, Marie Baekvad-Hansen, Steven Bakker, Gavin Band, Jack D Barchas, Ines Barroso, Nicholas Bass, Michael Bauer, Bernhard T Baune, Martin Begemann, Celine Bellenguez, Richard A Belliveau, Frank Bellivier, Stephan Bender, Judit Bene, Sarah E Bergen, Wade H Berrettini, Elizabeth Bevilacqua, Joanna M Biernacka, Tim B Bigdeli, Donald W Black, Hannah Blackburn, Jenefer M Blackwell, Douglas HR Blackwood, Carsten Bocker Pedersen, Michael Boehnke, Marco Boks, Anders D Borglum, Elvira Bramon, Gerome Breen, Matthew A Brown, Richard Bruggeman, Nancy G Buccola, Randy L Buckner, Monika Budde, Brendan Bulik-Sullivan, Suzannah J Bumpstead, William Bunney, Margit Burmeister, Joseph D Buxbaum, Jonas Bybjerg-Grauholm, William Byerley, Wiepke Cahn, Guiqing Cai, Murray J Cairns, Dominique Campion, Rita M Cantor, Vaughan J Carr, Noa Carrera, Juan P Casas, Miquel Casas, Stanley V Catts, Pablo Cervantes, Kimberley D Chambert, Raymond CK Chan, Eric YH Chen, Ronald YL Chen, Wei Cheng, Eric FC Cheung, Siow Ann Chong, Toni-Kim Clarke, C Robert Cloninger, David Cohen, Nadine Cohen, Jonathan R I Coleman, David A Collier, Paul Cormican, William Coryell, Nicholas Craddock, David W Craig, Benedicto Crespo-Facorro, James J Crowley, Cristiana Cruceanu, David Curtis, Piotr M Czerski, Anders M Dale, Mark J Daly, Udo Dannlowski, Ariel Darvasi, Michael Davidson, Kenneth L Davis, Christiaan A de Leeuw, Franziska Degenhardt, Jurgen Del Favero, Lynn E DeLisi, Panos Deloukas, Ditte Demontis, J Raymond DePaulo, Marta di Forti, Dimitris Dikeos, Timothy Dinan, Srdjan Djurovic, Amanda L Dobbyn, Peter Donnelly, Gary Donohoe, Elodie Drapeau, Serge Dronov, Jubao Duan, Frank Dudbridge, Audrey Duncanson, Howard Edenberg, Sarah Edkins, Hannelore Ehrenreich, Peter Eichhammer, Torbjorn Elvsashagen, Johan Eriksson, Valentina Escott-Price, Tonu Esko, Laurent Essioux, Bruno Etain, Chun Chieh Fan, Kai-How Farh, Martilias S Farrell, Matthew Flickinger, Tatiana M Foroud, Liz Forty, Josef Frank, Lude Franke, Christine Fraser, Robert Freedman, Colin Freeman, Nelson B Freimer, Joseph I Friedman, Menachem Fromer, Mark A Frye, Janice M Fullerton, Katrin Gade, Julie Garnham, Helena A Gaspar, Pablo V Gejman, Giulio Genovese, Lyudmila Georgieva, Claudia Giambartolomei, Eleni Giannoulatou, Ina Giegling, Michael Gill, Matthew Gillman, Marianne Giortz Pedersen, Paola Giusti-Rodriguez, Stephanie Godard, Fernando Goes, Jacqueline I Goldstein, Srihari Gopal, Scott D Gordon, Katherine Gordon-Smith, Jacob Gratten, Emma Gray, Elaine K Green, Melissa J Green, Tiffany A Greenwood, Maria Grigoroiu-Serbanescu, Jakob Grove, Weihua Guan, Hugh Gurling, Jose Guzman Parra, Rhian Gwilliam, Lieuwe de Haan, Jeremy Hall, Mei-Hua Hall, Christian Hammer, Naomi Hammond, Marian L Hamshere, Mark Hansen, Thomas Hansen, Vahram Haroutunian, Annette M Hartmann, Joanna Hauser, Martin Hautzinger, Urs Heilbronner, Garrett Hellenthal, Frans A Henskens, Stefan Herms, Maria Hipolito, Joel N Hirschhorn, Per Hoffmann, Mads V Hollegaard, David M Hougaard, Hailiang Huang, Laura Huckins, Christina M Hultman, Sarah E Hunt, Masashi Ikeda, Nakao Iwata, Conrad Iyegbe, Assen V Jablensky, Stephane Jamain, Janusz Jankowski, Alagurevathi Jayakumar, Inge Joa, Ian Jones, Lisa A Jones, Erik G Jonsson, Antonio Julia, Anders Jureus, Anna K Kahler, Rene S Kahn, Luba Kalaydjieva, Radhika Kandaswamy, Sena Karachanak-Yankova, Juha Karjalainen, Robert Karlsson, David Kavanagh, Matthew C Keller, Brian J Kelly, John Kelsoe, James L Kennedy, Andrey Khrunin, Yunjung Kim, George Kirov, Sarah Kittel-Schneider, Janis Klovins, Jo Knight, Sarah V Knott, James A Knowles, Manolis Kogevinas, Bettina Konte, Eugenia Kravariti, Vaidutis Kucinskas, Zita Ausrele Kucinskiene, Ralph Kupka, Hana Kuzelova-Ptackova, Mikael Landen, Cordelia Langford, Claudine Laurent, Jacob Lawrence, Stephen Lawrie, William B Lawson, Markus Leber, Marion Leboyer, Phil H Lee, Jimmy Lee Chee Keong, Sophie E Legge, Todd Lencz, Bernard Lerer, Douglas F Levinson, Shawn E Levy, Cathryn M Lewis, Jun Z Li, Miaoxin Li, Qingqin S Li, Tao Li, Kung-Yee Liang, Jennifer Liddle, Jeffrey Lieberman, Svetlana Limborska, Kuang Lin, Don H Linszen, Jolanta Lissowska, Chunyu Liu, Jianjun Liu, Jouko Lonnqvist, Carmel M Loughland, Jan Lubinski, Susanne Lucae, Milan Macek, Donald J MacIntyre, Patrik KE Magnusson, Brion S Maher, Pamela B Mahon, Wolfgang Maier, Anil K Malhotra, Jacques Mallet, Ulrik F Malt, Hugh S Markus, Sara Marsal, Nicholas G Martin, Ignacio Mata, Christopher G Mathew, Manuel Mattheisen, Morten Mattingsdal, Fermin Mayoral, Owen T McCann, Robert W McCarley, Steven A McCarroll, Mark I McCarthy, Colm McDonald, Susan L McElroy, Peter McGuffin, Melvin G Mclnnis, Andrew M McIntosh, James D McKay, Francis J McMahon, Helena Medeiros, Sarah E Medland, Sandra Meier, Carin J Meijer, Bela Melegh, Ingrid Melle, Fan Meng, Raquelle I Mesholam-Gately, Andres Metspalu, Patricia T Michie, Lili Milani, Vihra Milanova, Philip B Mitchell, Younes Mokrab, Grant W Montgomery, Jennifer L Moran, Gunnar Morken, Derek W Morris, Ole Mors, Preben B Mortensen, Bryan J Mowry, Thomas W Mühleisen, Bertram Müller-Myhsok, Kieran C Murphy, Robin M Murray, Richard M Myers, Inez Myin-Germeys, Benjamin M Neale, Mari Nelis, Igor Nenadic, Deborah A Nertney, Gerald Nestadt, Kristin K Nicodemus, Caroline M Nievergelt, Liene Nikitina-Zake, Vishwajit Nimgaonkar, Laura Nisenbaum, Merete Nordentoft, Annelie Nordin, Markus M Nöthen, Evaristus A Nwulia, Eadbhard O’Callaghan, Claire O’Donovan, O’Dushlaine Colm, F Anthony O’Neill, Ketil J Oedegaard, Sang-Yun Oh, Ann Olincy, Line Olsen, Lilijana Oruc, Jim Van Os, Michael J Owen, Sara A Paciga, Colin N A Palmer, Aarno Palotie, Christos Pantelis, George N Papadimitriou, Elena Parkhomenko, Carlos Pato, Michele T Pato, Tiina Paunio, Richard Pearson, Psychosis Endophenotypes International Consortium, Diana O Perkins, Roy H Perlis, Amy Perry, Tune H Pers, Tracey L Petryshen, Andrea Pfennig, Marco Picchioni, Olli Pietilainen, Jonathan Pimm, Matti Pirinen, Robert Plomin, Andrew J Pocklington, Danielle Posthuma, James B Potash, Simon C Potter, John Powell, Alkes Price, Ann E Pulver, Shaun M Purcell, Digby Quested, Josep Antoni Ramos-Quiroga, Henrik B Rasmussen, Anna Rautanen, Radhi Ravindrarajah, Eline J Regeer, Abraham Reichenberg, Andreas Reif, Mark A Reimers, Marta Ribases, John P Rice, Alexander L Richards, Michelle Ricketts, Brien P Riley, Fabio Rivas, Margarita Rivera, Joshua L Roffman, Guy A Rouleau, Panos Roussos, Dan Rujescu, Veikko Salomaa, Cristina Sanchez-Mora, Alan R Sanders, Stephen J Sawcer, Ulrich Schall, Alan F Schatzberg, William A Scheftner, Peter R Schofield, Nicholas J Schork, Sibylle G Schwab, Edward M Scolnick, Laura J Scott, Rodney J Scott, Larry J Seidman, Alessandro Serretti, Pak C Sham, Cynthia Shannon Weickert, Tatyana Shehktman, Jianxin Shi, Paul D Shilling, Engilbert Sigurdsson, Jeremy M Silverman, Kang Sim, Claire Slaney, Petr Slominsky, Olav B Smeland, Jordan W Smoller, Hon-Cheong So, Janet L Sobell, Erik Soderman, Christine Soholm Hansen, Chris C A Spencer, Anne T Spijker, David St Clair, Hreinn Stefansson, Kari Stefansson, Stacy Steinberg, Elisabeth Stogmann, Eystein Stordal, Amy Strange, Richard E Straub, John S Strauss, Fabian Streit, Eric Strengman, Jana Strohmaier, T Scott Stroup, Zhan Su, Mythily Subramaniam, Jaana Suvisaari, Dragan M Svrakic, Jin P Szatkiewicz, Szabolcs Szelinger, Avazeh Tashakkori-Ghanbaria, Srinivas Thirumalai, Robert C Thompson, Thorgeir E Thorgeirsson, Draga Toncheva, Paul A Tooney, Sarah Tosato, Timothea Toulopoulou, Richard C Trembath, Jens Treutlein, Vassily Trubetskoy, Gustavo Turecki, Arne E Vaaler, Helmut Vedder, Eduard Vieta, John Vincent, Peter M Visscher, Ananth C Viswanathan, Damjan Vukcevic, John Waddington, Matthew Waller, Dermot Walsh, Muriel Walshe, James TR Walters, Dai Wang, Qiang Wang, Weiqing Wang, Yunpeng Wang, Stanley J Watson, Bradley T Webb, Thomas W Weickert, Daniel R Weinberger, Matthias Weisbrod, Mark Weiser, Thomas Werge, Paul Weston, Pamela Whittaker, Sara Widaa, Durk Wiersma, Dieter B Wildenauer, Nigel M Williams, Stephanie Williams, Stephanie H Witt, Aaron R Wolen, Emily HM Wong, Nicholas W Wood, Brandon K Wormley, Wellcome Trust Case-Control Consortium, Jing Qin Wu, Simon Xi, Wei Xu, Allan H Young, Clement C Zai, Peter Zandi, Peng Zhang, Xuebin Zheng, Fritz Zimprich, Sebastian Zollner, Aiden Corvin, Ayman H Fanous, Sven Cichon, Marcella Rietschel, Elliot S Gershon, Thomas G Schulze, Alfredo B Cuellar-Barboza, Andreas J Forstner, Peter A Holmans, John I Nurnberger, Ole A Andreassen, S Hong Lee, Michael C O’Donovan, Patrick F Sullivan, Roel A Ophoff, Naomi R Wray, Pamela Sklar, Kenneth S Kendler

**Affiliations:** Division of Genetic Medicine, Departments of Medicine, Psychiatry, Biomedical Informatics, Vanderbilt Genetics Institute, Vanderbilt University Medical Center, Nashville, TN, USA; Analytic and Translational Genetics Unit, Massachusetts General Hospital, Boston, Massachusetts, USA; Department of Psychiatry and Psychotherapy, Charite - Universitatsmedizin, Berlin, Germany; Stanley Center for Psychiatric Research, Broad Institute of MIT and Harvard, Cambridge, Massachusetts, USA; Molecular Psychiatry Laboratory, Division of Psychiatry, University College London, London, UK; Department of Human Genetics, David Geffen School of Medicine, University of California, Los Angeles, California, USA; Department of Genetics and Genomic Sciences, Icahn School of Medicine at Mount Sinai, New York, NY USA; Queensland Brain Institute, The University of Queensland, Brisbane, Australia; MRC Social, Genetic and Developmental Psychiatry Centre, King’s College London, London, UK; Department of Psychiatry, Icahn School of Medicine at Mount Sinai, New York, New York, USA; Center for Neurobehavioral Genetics, Semel Institute for Neuroscience and Human Behavior, University of California, Los Angeles, California, USA; Centre for Integrative Biology, University of Trento, Trento, Italy; MRC Centre for Neuropsychiatric Genetics and Genomics, Division of Psychological Medicine and Clinical Neurosciences, Cardiff University, Cardiff, UK; Institute of Psychiatric Phenomics and Genomics (IPPG), Medical Center of the University of Munich, Munich, Germany; Department of Psychiatry and Psychotherapy, Ludwig Maximillian University, Munich, Germany; Department of Clinical Sciences, Psychiatry, Umea University, Umea, Sweden; Department of Clinical Neuroscience, Psychiatry Section, Karolinska Institutet, Stockholm, Sweden; Department of Psychiatry, Diakonhjemmet Hospital, Oslo, Norway; NORMENT, KG Jebsen Centre for Psychosis Research, Institute of Clinical Medicine, University of Oslo, Oslo, Norway; Centre for Integrative Register-based Research, CIRRAU, Aarhus University, Aarhus, Denmark; National Centre for Register-based Research, Aarhus University, Aarhus, Denmark; The Lundbeck Foundation Initiative for Integrative Psychiatric Research, iPSYCH, Denmark; Molecular & Behavioral Neuroscience Institute, University of Michigan, Ann Arbor, MI USA; NEUROSCIENCE, Istituto Di Ricerche Farmacologiche Mario Negri, Milano, Italy; State Mental Hospital, Haar, Germany; Department of Psychiatry, Dalhousie University, Halifax, NS Canada; National Institute of Mental Health, Klecany, Czech Republic; Department of Psychiatry and Behavioral Sciences, Stanford University, Stanford, California, USA; Medical Research Council Centre for Neuropsychiatric Genetics and Genomics, Division of Psychological Medicine and Clinical Neurosciences, Cardiff University, Cardiff, UK; Department of Psychiatry and Behavioral Neuroscience, University of Chicago, Chicago, Illinois, USA; Department of Biomedicine, Aarhus University, Aarhus, Denmark; iSEQ, Centre for Integrative Sequencing, Aarhus University, Aarhus, Denmark; Department of Psychiatry and Behavioral Sciences, Atlanta Veterans Affairs Medical Center, Atlanta, Georgia, USA; Department of Psychiatry and Behavioral Sciences, Emory University, Atlanta, Georgia, USA; Psychiatry, Berkshire Healthcare NHS Foundation Trust, Bracknell, UK; Fundacio de Docencia i Recerca Mutua de Terrassa, Universitat de Barcelona, Spain; Institute of Psychiatry at King’s College London, London, UK; Virginia Institute for Psychiatric and Behavioral Genetics, Department of Psychiatry, Virginia Commonwealth University, Richmond, Virginia, USA; Psychiatry, Rush University Medical Center, Chicago, IL USA; Statens Serum Institut, Copenhagen, Denmark; Department of Psychiatry, Rudolf Magnus Institute of Neuroscience, University Medical Center Utrecht, Utrecht, The Netherlands; Wellcome Trust Centre for Human Genetics, Oxford, UK; Department of Psychiatry, Weill Cornell Medical College, New York, NY USA; Wellcome Trust Sanger Institute, Wellcome Trust Genome Campus, Hinxton, Cambridge, UK; Division of Psychiatry, University College London, London, GB; Department of Psychiatry and Psychotherapy, University Hospital Carl Gustav Carus, Technische Universitat Dresden, Dresden, Germany; Discipline of Psychiatry, University of Adelaide, Adelaide, Australia; Clinical Neuroscience, Max Planck Institute of Experimental Medicine, Gottingen, Germany; Department of Psychiatry and Addiction Medicine, Assistance Publique - Hopitaux de Paris, Paris, France; Paris Bipolar and TRD Expert Centres, FondaMental Foundation, Paris, France; Psychiatry, Universite Paris Diderot, Paris, France; UMR-S1144 Team 1: Biomarkers of relapse and therapeutic response in addiction and mood disorders, INSERM, Paris, France; Child and Adolescent Psychiatry, University of Technology Dresden, Dresden, Germany; Section for Experimental Psychopathology, General Psychiatry, Heidelberg, Germany; Department of Medical Genetics, University of Pecs, Pecs, Hungary; Szentagothai Research Center, University of Pecs, Pecs, Hungary; Department of Medical Epidemiology and Biostatistics, Karolinska Institutet, Stockholm, Sweden; Psychiatry, University of Pennsylvania, Philadelphia, PA USA; Health Sciences Research, Mayo Clinic, Rochester, Minnesota USA; Department of Psychiatry, University of Iowa Carver College of Medicine, Iowa City, Iowa, USA; Cambridge Institute for Medical Research, University of Cambridge School of Clinical Medicine, Cambridge, UK; Telethon Institute for Child Health Research, Centre for Child Health Research, University of Western Australia, subiaco, Western Australia, Australia; Division of Psychiatry, University of Edinburgh, Edinburgh, UK; Center for Statistical Genetics and Department of Biostatistics, University of Michigan, Ann Arbor, Michigan USA; Department of Biomedicine, Aarhus University, Aarhus C, Denmark; Institute of Cognitive Neuroscience, University College London, London, UK; Mental Health Sciences Unit, University College London, London, UK; NIHR BRC for Mental Health, King’s College London, London, UK; Diamantina Institute of Cancer, Immunology and Metabolic Medicine, Princess Alexandra Hospital, University of Queensland, Brisbane, Queensland, Australia; University Medical Center Groningen, Department of Psychiatry, University of Groningen, RB, The Netherlands; School of Nursing, Louisiana State University Health Sciences Center, New Orleans, Louisiana, USA; Athinoula A. Martinos Center, Massachusetts General Hospital, Boston, Massachusetts, USA; Center for Brain Science, Harvard University, Cambridge, Massachusetts, USA; Department of Psychiatry, Massachusetts General Hospital, Boston, Massachusetts, USA; Department of Psychiatry and Human Behavior, University of California, Irvine, Irvine, California USA; Molecular & Behavioral Neuroscience Institute and Department of Computational Medicine & Bioinformatics, University of Michigan, Ann Arbor, Michigan USA; Department of Human Genetics, Icahn School of Medicine at Mount Sinai, New York, New York, USA; Department of Neuroscience, Icahn School of Medicine at Mount Sinai, New York, New York, USA; Friedman Brain Institute, Icahn School of Medicine at Mount Sinai, New York, New York, USA; Neonatal Genetik, Statens Serum Institut, Copenhagen, Denmark; Department of Psychiatry, University of California San Francisco, San Francisco, California, USA; Priority Centre for Translational Neuroscience and Mental Health, University of Newcastle, Newcastle, Australia; Schizophrenia Research Institute, Sydney, Australia; School of Biomedical Sciences and Pharmacy, University of Newcastle, Callaghan, Australia; Centre Hospitalier du Rouvray and INSERM U 1079Faculty of Medicine, Rouen, France; School of Psychiatry, University of New South Wales, Sydney, Australia; MRC Centre for Neuropsychiatric Genetics and Genomics, Institute of Psychological Medicine and Clinical Neurosciences, School of Medicine, Cardiff University, Cardiff, UK; Department of Epidemiology and Population Health, London School of Hygiene and Tropical Medicine, London, UK; Department of Epidemiology and Public Health, University College London, London, UK; Department of Psychiatry and Forensic Medicine, Universitat Autonoma de Barcelona, Barcelona, Spain; Department of Psychiatry, Hospital Universitari Vall ďHebron, Barcelona, Spain; Instituto de Salud Carlos III, Biomedical Network Research Centre on Mental Health (CIBERSAM), Madrid, Spain; Psychiatric Genetics Unit, Group of Psychiatry Mental Health and Addictions, Vall ďHebron Research Institut (VHIR), Universitat Autonoma de Barcelona, Barcelona, Spain; Royal Brisbane and Women’s Hospital, University of Queensland, Brisbane, Australia; Department of Psychiatry, Mood Disorders Program, McGill University Health Center, Montreal, QC Canada; Institute of Psychology, Chinese Academy of Science, Beijing, China; Department of Psychiatry, Li Ka Shing Faculty of Medicine, The University of Hong Kong, Hong Kong, China; State Key Laboratory for Brain and Cognitive Sciences, Li Ka Shing Faculty of Medicine, The University of Hong Kong, Hong Kong, China; Department of Computer Science, University of North Carolina, Chapel Hill, North Carolina, USA; Castle Peak Hospital, Hong Kong, China; Institute of Mental Health, Singapore, Singapore; Department of Psychiatry, Washington University, St. Louis, Missouri, USA; Department of Child and Adolescent Psychiatry, Assistance Publique Hospitaux de Paris, Pierre and Marie Curie Faculty of Medicine and Institute for Intelligent Systems and Robotics, Paris, France; Blue Note Biosciences, Princeton, New Jersey, USA; Eli Lilly and Company Limited, Erl Wood Manor, Sunninghill Road, Windlesham, Surrey, UK; Social, Genetic and Developmental Psychiatry Centre, Institute of Psychiatry, King’s College London, London, UK; Neuropsychiatric Genetics Research Group, Department of Psychiatry, Trinity College Dublin, Ireland; University of Iowa Hospitals and Clinics, Iowa City, Iowa, USA; National Centre for Mental Health, Cardiff University, Cardiff, UK; Translational Genomics, USAC, Phoenix, Arizona, USA; Centro Investigacion Biomedica en Red Salud Mental, Madrid, Spain; University Hospital Marques de Valdecilla, Instituto de Formacion e Investigacion Marques de Valdecilla, University of Cantabria, Santander, Spain; Department of Genetics, University of North Carolina, Chapel Hill, North Carolina, USA; Department of Translational Research in Psychiatry, Max Planck Institute of Psychiatry, Munich, Germany; Centre for Psychiatry, Queen Mary University of London, London, UK; UCL Genetics Institute, University College London, London, UK; Department of Psychiatry, Laboratory of Psychiatric Genetics, Poznan University of Medical Sciences, Poznan, Poland; Neurosciences, Radiology, Psychiatry, Cognitive Science, University of California San Diego, La Jolla, California USA; Medical and Population Genetics Program, Broad Institute of MIT and Harvard, Cambridge, Massachusetts, USA; Department of Psychiatry, University of Munster, Munster, Germany; Department of Genetics, The Hebrew University of Jer USA lem, JerUSAlem, Israel; Sheba Medical Center, Tel Hashomer, Israel; Department of Complex Trait Genetics, Center for Neurogenomics and Cognitive Research Neuroscience, Vrije Universiteit Amsterdam, Amsterdam, The Netherlands; Life&Brain Center, Department of Genomics, University of Bonn, Bonn, Germany; Institute of Human Genetics, University of Bonn, Bonn, Germany; Applied Molecular Genomics Unit, VIB Department of Molecular Genetics, University of Antwerp, Antwerp, Belgium; Department of Psychiatry, Harvard Medical School, Boston, Massachusetts, USA; VA Boston Health Care System, Brockton, Massachusetts, USA; Department of Psychiatry and Behavioral Sciences, Johns Hopkins University School of Medicine, Baltimore, Maryland, USA; King’s College London, London, UK; First Department of Psychiatry, University of Athens Medical School, Athens, Greece; Department of Psychiatry, University College Cork, Co. Cork, Ireland; Department of Medical Genetics, Oslo University Hospital, Oslo, Norway; Division of Psychiatric Genomics, Department of Psychiatry, Icahn School of Medicine at Mount Sinai, New York, New York, USA; Department of Statistics, University of Oxford, Oxford, UK; Cognitive Genetics and Therapy Group, School of Psychology and Discipline of Biochemistry, National University of Ireland Galway, Co. Galway, Ireland; Department of Psychiatry and Behavioral Sciences, NorthShore University HealthSystem, Evanston, Illinois, USA; Department of Non-Communicable Disease Epidemiology, London School of Hygiene and Tropical Medicine, London, UK; Molecular and Physiological Sciences, The Wellcome Trust, London, UK; Biochemistry and Molecular Biology, Indiana University School of Medicine, Indianapolis, Indiana USA; Department of Psychiatry, University of Regensburg, Regensburg, Germany; Department of Neurology, Oslo University Hospital, Oslo, Norway; Department of General Practice, Helsinki University Central Hospital, University of Helsinki, Helsinki, Finland; Folkhalsan Research Center, Helsinki, Finland, Biomedicum Helsinki 1, Haartmaninkatu 8, Helsinki, Finland; National Institute for Health and Welfare, Helsinki, Finland; Department of Genetics, Harvard Medical School, Boston, Massachusetts, USA; Division of Endocrinology and Center for Basic and Translational Obesity Research, Boston Children’s Hospital, Boston, Massachusetts, USA; Estonian Genome Center, University of Tartu, Tartu, Estonia; Translational Technologies and Bioinformatics, Pharma Research and Early Development, F.Hoffman-La Roche, Basel, Switzerland; Centre for Affective Disorders, Institute of Psychiatry, Psychology and Neuroscience, London, UK; Cognitive Science, University of California San Diego, La Jolla, California USA; Department of Medical & Molecular Genetics, Indiana University, Indianapolis, Indiana USA; Department of Genetic Epidemiology in Psychiatry, Central Institute of Mental Health, Medical Faculty Mannheim, University of Heidelberg, Heidelberg, Mannheim, Germany; Department of Genetics, University of Groningen, University Medical Centre Groningen, Groningen, The Netherlands; Department of Psychiatry, University of Colorado Denver, Aurora, Colorado, USA; Psychiatric and Neurodevelopmental Genetics Unit, Massachusetts General Hospital, Boston, Massachusetts, USA; Department of Psychiatry & Psychology, Mayo Clinic, Rochester, Minnesota USA; Neuroscience Research Australia, Sydney, Australia; School of Medical Sciences, University of New South Wales, Sydney, Australia; Department of Psychiatry and Psychotherapy, University Medical Center Gottingen, Gottingen, Germany; Department of Psychiatry, University of Halle, Halle, Germany; Department of Psychiatry, University of Munich, Munich, Germany; Neuropsychiatric Genetics Research Group, Dept of Psychiatry and Trinity Translational Medicine Institute, Trinity College Dublin, Dublin, Ireland; Department of Economics and Business Economics, National Centre for Register-based Research, Aarhus University, Aarhus, Denmark; Departments of Psychiatry and Human and Molecular Genetics, INSERM, Institut de Myologie, Hopital de la Pitie-Salpetriere, Paris, France; Neuroscience Therapeutic Area, Janssen Research and Development, Raritan, New Jersey, USA; Genetics and Computational Biology, QIMR Berghofer Medical Research Institute, Brisbane, Australia; Department of Psychological Medicine, University of Worcester, Worcester, UK; School of Biomedical and Healthcare Sciences, Plymouth University Peninsula Schools of Medicine and Dentistry, Plymouth, UK; Department of Psychiatry, University of California San Diego, La Jolla, California USA; Biometric Psychiatric Genetics Research Unit, Alexandru Obregia Clinical Psychiatric Hospital, Bucharest, Romania; Bioinformatics Research Centre (BiRC), Aarhus University, Aarhus, Denmark; Biostatistics, University of Minnesota System, Minneapolis, MN USA; Mental Health Department, University Regional Hospital. Biomedicine Institute (IBIMA), Malaga, Spain; Academic Medical Centre University of Amsterdam, Department of Psychiatry, Amsterdam, The Netherlands; Mclean Hospital, Harvard Medical School, 115 Mill St, Belmont Massachusetts, USA; Illumina, La Jolla, California, California, USA; Institute of Biological Psychiatry, Mental Health Centre Sct. Hans, Mental Health Services Copenhagen, Denmark; J.J. Peters VA Medical Center, Bronx, New York, New York, USA; Department of Psychology, Eberhard Karls Universitat Tobingen, Tubingen, Germany; Priority Centre for Health Behaviour, University of Newcastle, Newcastle, Australia; School of Electrical Engineering and Computer Science, University of Newcastle, Newcastle, Australia; Department of Biomedicine, University of Basel, Basel, Switzerland; Department of Psychiatry and Behavioral Sciences, Howard University Hospital, Washington, DC USA; Institute of Medical Genetics and Pathology, University Hospital Basel, Basel, Switzerland; Section of Neonatal Screening and Hormones, Department of Clinical Biochemistry, Immunology and Genetics, Statens Serum Institut, Copenhagen, Denmark; Department for Congenital Disorders, Statens Serum Institut, Copenhagen, Denmark; Department of Psychiatry, Fujita Health University School of Medicine, Toyoake, Aichi, Japan; Centre for Clinical Research in Neuropsychiatry, School of Psychiatry and Clinical Neurosciences, The University of Western Australia, Medical Research Foundation Building, Perth, Australia; School of Psychiatry and Clinical Neurosciences, The University of Western Australia, Perth, Australia; The Perkins Institute for Medical Research,The University of Western Australia, Perth, Australia; Faculte de Medecine, Universite Paris Est, Creteil, France; Psychiatrie Translationnelle, Inserm U955, Creteil, France; Peninsula School of Medicine and Dentistry, Plymouth University, Plymouth, UK; Regional Centre for Clinical Research in Psychosis, Department of Psychiatry, Stavanger University Hospital, Stavanger, Norway; Rheumatology Research Group, Vall ďHebron Research Institute, Barcelona, Spain; Centre for Medical Research, The University of Western Australia, Perth, Western Australia, Australia; Western Australian Institute for Medical Research, The University of Western Australia, Perth, Western Australia, Australia; Department of Medical Genetics, Medical University, Sofia, Bulgaria; Department of Psychology, University of Colorado Boulder, Boulder, Colorado, USA; Campbell Family Mental Health Research Institute, Centre for Addiction and Mental Health, Toronto, Ontario, Canada; Department of Psychiatry, University of Toronto, Toronto, Ontario, Canada; Institute of Medical Science, University of Toronto, Toronto, Ontario, Canada; Institute of Molecular Genetics, Russian Academy of Sciences, Moscow, Russia; Department of Psychiatry, Psychosomatic Medicine and Psychotherapy, University Hospital Frankfurt, Frankfurt am Main, Germany; Latvian Biomedical Research and Study Centre, Riga, Latvia; Cell Biology, SUNY Downstate Medical Center College of Medicine, Brooklyn, NY USA; Department of Psychiatry and Zilkha Neurogenetics Institute, Keck School of Medicine at University of Southern California, Los Angeles, California, USA; Institute for Genomic Health, SUNY Downstate Medical Center College of Medicine, Brooklyn, NY USA; Center for Research in Environmental Epidemiology (CREAL), Barcelona, Spain; Faculty of Medicine, Vilnius University, Vilnius, Lithuania; Psychiatry, GGZ inGeest, Amsterdam, The Netherlands; Psychiatry, VU medisch centrum, Amsterdam, The Netherlands; Department of Biology and Medical Genetics, 2nd Faculty of Medicine and University Hospital Motol, Prague, Czech Republic; Institute of Neuroscience and Physiology, University of Gothenburg, Gothenburg, Sweden; Department of Child and Adolescent Psychiatry, Pierre and Marie Curie Faculty of Medicine, Paris, France; Psychiatry, North East London NHS Foundation Trust, Ilford, UK; Clinic for Psychiatry and Psychotherapy, University Hospital Cologne, Cologne, Germany; INSERM, Paris, France; Duke-NUSA Graduate Medical School, Singapore, Singapore; Hofstra Northwell School of Medicine, Hempstead, New York, USA; The Feinstein Institute for Medical Research, Manhasset, New York, USA; The Hofstra NS-LIJ School of Medicine, Hempstead, New York, USA; Department of Psychiatry, Hadassah-Hebrew University Medical Center, Jerusalem, Israel; HudsonAlpha Institute for Biotechnology, Huntsville, AL USA; Department of Medical & Molecular Genetics, King’s College London, London, UK; Department of Human Genetics, University of Michigan, Ann Arbor, MI USA; Centre for Genomic Sciences, The University of Hong Kong, Hong Kong, China; Mental Health Centre and Psychiatric Laboratory, West China Hospital, Sichuan University, Chengdu, Sichuan, China; Department of Biostatistics, Johns Hopkins University Bloomberg School of Public Health, Baltimore, Maryland, USA; Department of Psychiatry, Columbia University, New York, New York, USA; Department of Psychiatry, Academic Medical Center, University of Amsterdam, Amsterdam, The Netherlands; Cancer Epidemiology and Prevention, M. Sklodowska-Curie Cancer Center and Institute of Oncology, Warsaw, Poland; Psychiatry, University of Illinois at Chicago College of Medicine, Chicago, IL USA; Human Genetics, Genome Institute of Singapore, A∗STAR, Singapore, Singapore; Saw Swee Hock School of Public Health, National University of Singapore, Singapore, Singapore; Department of Mental Health and substance Abuse Services; National Institute for Health and Welfare, Helsinki, Finland; Department of Genetics and Pathology, International Hereditary Cancer Center, Pomeranian Medical University in Szczecin, Szczecin, Poland; Max Planck Institute of Psychiatry, Munich, Germany; Division of Psychiatry, Centre for Clinical Brain Sciences, University of Edinburgh, Edinburgh, UK; Mental Health, NHS 24, Glasgow, UK; Department of Mental Health, Bloomberg School of Public Health, Johns Hopkins University, Baltimore, Maryland, USA; Psychiatry, Brigham and Women’s Hospital, Boston, MA USA; Department of Psychiatry and Psychotherapy, University of Bonn, Bonn, Germany; The Zucker Hillside Hospital, Glen Oaks, New York, USA; Centre National de la Recherche Scientifique, Laboratoire de Genetique Moleculaire de la Neurotransmission et des Processus Neurodegeneratifs, Hopital de la Pitie Salpetriere, Paris, France; Research and Education, Division of Clinical Neuroscience, Oslo Universitetssykehus, Oslo, Norway; Clinical Neurosciences, St George’s University of London, London, UK; School of Psychology, The University of Queensland, Brisbane, Australia; Department of Medical and Molecular Genetics, School of Medicine, King’s College London, Guy’s Hospital, London, UK; Department of Genomics Mathematics, University of Bonn, D-53127 Bonn, Germany; Stockholm Health Care Services, Stockholm County Council, Stockholm, Sweden; Research Unit, Sorlandet Hospital, Kristiansand, Norway; Oxford Centre for Diabetes, Endocrinology and Metabolism, Churchill Hospital, Oxford, UK; Department of Psychiatry, National University of Ireland Galway, Co. Galway, Ireland; Research Institute, Lindner Center of HOPE, Mason, OH USA; Department of Psychiatry, University of Michigan, Ann Arbor, MI USA; Centre for Cognitive Ageing and Cognitive Epidemiology, University of Edinburgh, Edinburgh, UK; Genetic Cancer Susceptibility Group, International Agency for Research on Cancer, Lyon, France; Human Genetics Branch, Intramural Research Program, National Institute of Mental Health, Bethesda, MD USA; Division of Mental Health and Addiction, Oslo University Hospital, Oslo, Norway; Massachusetts Mental Health Center Public Psychiatry Division of the Beth Israel Deaconess Medical Center, Boston, Massachusetts, USA; Institute of Molecular and Cell Biology, University of Tartu, Tartu, Estonia; School of Psychology, University of Newcastle, Newcastle, Australia; First Psychiatric Clinic, Medical University, Sofia, Bulgaria; Institute for Molecular Bioscience, The University of Queensland, Brisbane, Australia; Mental Health, Faculty of Medicine and Health Sciences, Norwegian University of Science and Technology - NTNU, Trondheim, Norway; Psychiatry, St Olavs University Hospital, Trondheim, Norway; Discipline of Biochemistry, Neuroimaging and Cognitive Genomics (NICOG) Centre, National University of Ireland, Galway, Galway, Ireland; Queensland Centre for Mental Health Research, University of Queensland, Brisbane, Australia; Institute of Neuroscience and Medicine (INM-1), Research Center Juelich, Juelich, Germany; Institute of Translational Medicine, University of Liverpool, Liverpool, UK; Munich Cluster for Systems Neurology (SyNergy), Munich, Germany; Department of Psychiatry, Royal College of Surgeons in Ireland, Dublin, Ireland; Maastricht University Medical Centre, South Limburg Mental Health Research and Teaching Network, EURON, Maastricht, The Netherlands; Department of Psychiatry and Psychotherapy, Jena University Hospital, Jena, Germany; Department of Psychiatry, Trinity College Dublin, Dublin, Ireland; Research/Psychiatry, Veterans Affairs San Diego Healthcare System, San Diego, California USA; Psychiatry and Human Genetics, University of Pittsburgh, Pittsburgh, PA USA; Eli Lilly and Company, Lilly Corporate Center, Indianapolis, Indiana, USA; Mental Health Centre Copenhagen, Copenhagen University Hospital, Copenhagen, Denmark; DETECT Early Intervention Service for Psychosis, Blackrock, Co. Dublin, Ireland; Centre for Public Health, Institute of Clinical Sciences, Queen’s University Belfast, Belfast, UK; Division of Psychiatry, Haukeland Universitetssjukehus, Bergen, Norway; Faculty of Medicine and Dentistry, University of Bergen, Bergen, Norway; Lawrence Berkeley National Laboratory, University of California at Berkeley, Berkeley, California, USA; Department of Clinical Psychiatry, Psychiatry Clinic, Clinical Center University of Sarajevo, Sarajevo, Bosnia; Human Genetics and Computational Biomedicine, Pfizer Global Research and Development, Groton, Connecticut, USA; Biomedical Research Centre, Ninewells Hospital and Medical School, Dundee, UK; Institute for Molecular Medicine Finland, FIMM, University of Helsinki, Helsinki, Finland; Melbourne Neuropsychiatry Centre, University of Melbourne & Melbourne Health, Melbourne, Australia; College of Medicine Institute for Genomic Health, SUNY Downstate Medical Center College of Medicine, Brooklyn, NY USA; Department of Psychiatry, University of Helsinki, Helsinki, Finland; Public Health Genomics Unit, National Institute for Health and Welfare, Helsinki, Finland; PEIC; Department of Psychiatry, University of North Carolina at Chapel Hill, Chapel Hill, North Carolina USA; Division of Clinical Research, Massachusetts General Hospital, Boston, MA USA; Center for Biological Sequence Analysis, Department of Systems Biology, Technical University of Denmark, Denmark; Center for Human Genetic Research and Department of Psychiatry, Massachusetts General Hospital, Boston, Massachusetts, USA; Department of Child and Adolescent Psychiatry, Erasmus University Medical Centre, Rotterdam, The Netherlands; Department of Complex Trait Genetics, Neuroscience Campus Amsterdam, VU University Medical Center Amsterdam, Amsterdam, The Netherlands; Department of Functional Genomics, Center for Neurogenomics and Cognitive Research, Neuroscience Campus Amsterdam, VU University, Amsterdam, The Netherlands; Department of Epidemiology, Harvard School of Public Health, Boston, Massachusetts, USA; Department of Psychiatry, University of Oxford, Oxford, UK; Outpatient Clinic for Bipolar Disorder, Altrecht, Utrecht, The Netherlands; Virginia Institute for Psychiatric and Behavioral Genetics, Virginia Commonwealth University, Richmond, Virginia, USA; Virginia Institute for Psychiatric and Behavioral Genetics, Departments of Psychiatry and Human and Molecular Genetics, Virginia Commonwealth University, Richmond, Virginia, USA; Department of Biochemistry and Molecular Biology II, Institute of Neurosciences, Center for Biomedical Research, University of Granada, Granada, Spain; Montreal Neurological Institute and Hospital, Montreal, QC Canada; Department of Neurology and Neurosurgery, McGill University, Faculty of Medicine, Montreal, QC Canada; Institute for Multiscale Biology, Icahn School of Medicine at Mount Sinai, New York, New York, USA; Department of Clinical Neurosciences, University of Cambridge, Addenbrooke’s Hospital, Cambridge, UK; Rush University Medical Center, Chicago, IL USA; Scripps Translational Science Institute, La Jolla, California USA; Faculty of Science, Medicine & Health, Univeristy of Wollogong, Australia; Hunter New England Health Service, Newcastle, Australia; Department of Biomedical and NeuroMotor Sciences, University of Bologna, Bologna, Italy; Division of Cancer Epidemiology and Genetics, National Cancer Institute, Bethesda, Maryland, USA; Faculty of Medicine, Department of Psychiatry, School of Health Sciences, University of Iceland, Reykjavik, Iceland; Research and Development, Bronx Veterans Affairs Medical Center, New York, New York, USA; Department of Neurosciences, University of California San Diego, La Jolla, California USA; Psychiatry and the Behavioral Sciences, University of Southern California, Los Angeles, California USA; Mood Disorders, PsyQ, Rotterdam, The Netherlands; University of Aberdeen, Institute of Medical Sciences, Aberdeen, UK; deCODE Genetics / Amgen, Reykjavik, Iceland; Faculty of Medicine, University of Iceland, Reykjavik, Iceland; Department of Clinical Neurology, Medical University of Vienna, Wien, Austria; Department of Neuroscience, Norges Teknisk Naturvitenskapelige Universitet Fakultet for naturvitenskap og teknologi, Trondheim, Norway; Department of Psychiatry, Hospital Namsos, Namsos, Norway; Lieber Institute for Brain Development, Baltimore, Maryland, USA; Centre for Addiction and Mental Health, Toronto, ON Canada; Department of Medical Genetics, University Medical Centre Utrecht, Universiteitsweg, Utrecht, The Netherlands; Neurogenomics, TGen, Los Angeles, AZ USA; Berkshire Healthcare NHS Foundation Trust, Bracknell, UK; Priority Research Centre for Translational Neuroscience and Mental Health, University of Newcastle, Newcastle, Australia; Section of Psychiatry, University of Verona, Verona, Italy; Department of Psychology, The Jockey Club Tower, 6/F, the Centennial Campus, Pokfulam Road, The University of Hong Kong, Hong Kong, China; Department of Psychiatry, McGill University, Montreal, QC Canada; Dept of Psychiatry, Sankt Olavs Hospital Universitetssykehuset i Trondheim, Trondheim, Norway; Psychiatry, Psychiatrisches Zentrum Nordbaden, Wiesloch, Germany; Clinical Institute of Neuroscience, Hospital Clinic, University of Barcelona, IDIBAPS, CIBERSAM, Barcelona, Spain; Institute of Ophthalmology, University College London, London, UK; National Institute for Health Research, Biomedical Research Centre at Moorfields Eye Hospital, National Health Service Foundation Trust, London, UK; Molecular and Cellular Therapeutics, Royal College of Surgeons in Ireland, Dublin, Ireland; Health Research Board, Dublin, Ireland; Institute of Clinical Medicine, University of Oslo, Oslo, Norway; Departments of Psychiatry, Neurology, Neuroscience and Institute of Genetic Medicine, Johns Hopkins School of Medicine, Baltimore, Maryland, USA; Department of Clinical Medicine, University of Copenhagen, Copenhagen, Denmark; Department of Psychiatry, University Medical Center Groningen, University of Groningen, The Netherlands; Department of Molecular Neuroscience, Institute of Neurology, London, UK; WTCCC2; Computational Sciences Center of Emphasis, Pfizer Global Research and Development, Cambridge, MA USA; Dalla Lana School of Public Health, University of Toronto, Toronto, ON Canada; Department of Biostatistics, Princess Margaret Cancer Centre, Toronto, ON Canada; Psychological Medicine, Institute of Psychiatry, Psychology & Neuroscience, King’s College London, London, UK; Department of Mental Health, Johns Hopkins University Bloomberg School of Public Health, Baltimore, MD USA; Institute of Genetic Medicine, Johns Hopkins University School of Medicine, Baltimore, MD USA; Department of Psychiatry, Institute for Genomic Health SUNY Downstate Medical Center Brooklyn, NY, USA; Departments of Psychiatry and Human Genetics, University of Chicago, Chicago, Illinois, USA; Universidad Autonoma de Nuevo Leon, Department of Psychiatry, Monterrey, Mexico; Department of Psychiatry, Mayo Clinic, MN, USA; Department of Psychiatry (UPK), University of Basel, Basel, Switzerland; Department of Psychiatry and Stark Neurosciences Research Institute, Indiana University School of Medicine, Indianapolis, IN USA; Virginia Institute for Psychiatric and Behavioral Genetics and Department of Psychiatry, Virginia Commonwealth University, Richmond, Virginia, USA

## Abstract

Schizophrenia (SCZ) and bipolar disorder (BD) are highly heritable disorders that share a significant proportion of common risk variation. Understanding the genetic factors underlying the specific symptoms of these disorders will be crucial for improving diagnosis, intervention and treatment. In case-control data consisting of 53,555 cases (20,129 BD, 33,426 SCZ) and 54,065 controls, we identified 114 genome-wide significant loci (GWS) when comparing all cases to controls, of which 41 represented novel findings. Two genome-wide significant loci were identified when comparing SCZ to BD and a third was found when directly incorporating functional information. Regional joint association identified a genomic region of overlapping association in BD and SCZ with disease-independent causal variants indicating a fourth region contributing to differences between these disorders. Regional SNP-heritability analyses demonstrated that the estimated heritability of BD based on the SCZ GWS regions was significantly higher than that based on the average genomic region (91 regions, p = 1.2×10^−6^) while the inverse was not significant (19 regions, p=0.89). Using our BD and SCZ GWAS we calculated polygenic risk scores and identified several significant correlations with: 1) SCZ subphenotypes: negative symptoms (SCZ, p=3.6×10^−6^) and manic symptoms (BD, p=2×10^−5^), 2) BD subphenotypes: psychotic features (SCZ p=1.2×10^−10^, BD p=5.3×10^−5^) and age of onset (SCZ p=7.9×10^−4^). Finally, we show that psychotic features in BD has significant SNP-heritability (h^2^ _snp_=0.15, SE=0.06), and a significant genetic correlation with SCZ (r_g_=0.34) in addition there is a significant sign test result between SCZ GWAS and a GWAS of BD cases contrasting those with and without psychotic features (p=0.0038, one-side binomial test). For the first time, we have identified specific loci pointing to a potential role of 4 genes (*DARS2*, *ARFGEF2*, *DCAKD* and *GATAD2A*) that distinguish between BD and SCZ, providing an opportunity to understand the biology contributing to clinical differences of these disorders. Our results provide the best evidence so far of genomic components distinguishing between BD and SCZ that contribute directly to specific symptom dimensions.

## Introduction

Bipolar disorder (BD) and schizophrenia (SCZ) are severe psychiatric disorders and among the leading causes of disability worldwide^1^. Both disorders have significant genetic components with heritability estimates ranging from 60-80%^2^. A genetic-epidemiological genetic study demonstrated a substantial overlap between these two disorders with a genetic correlation from common variation near 0.6-0.7 and high relative risks (RR) among relatives of both BD and SCZ patients (RRs for parent/offspring: BD/BD: 6.4, BD/SCZ: 2.4; SCZ/BD: 5.2, SCZ/SCZ: 9.9)^3^. Despite shared genetics and symptomology, the current diagnostic systems^4,5^ represent BD and SCZ as distinct categorical entities differentiated on the basis of their clinical presentation, with BD characterized by predominant mood symptoms, mood-congruent delusions and an episodic disease course and SCZ considered a prototypical psychotic disorder. Further, premorbid cognitive impairment and reduced intelligence are more frequent and severe in SCZ than BD^6^. The genetic contributors to these phenotypic distinctions have yet to be elucidated and could aid in understanding the underlying biology of their unique clinical presentation.

While the shared genetic component is large, studies to date have identified key genetic architecture differences between these two disorders. A polygenic risk score created from a case only SCZ vs BD genome-wide association study (GWAS) significantly correlated with SCZ vs BD diagnosis in an independent sample^7^, providing evidence that differences between the disorders also have a genetic basis. An enrichment of rare, moderate to highly penetrant copy number variants (CNVs) and *de novo* CNVs are seen in SCZ patients^8–12^, while, the involvement of CNVs in BD is much less clear^13^. Although the role of *de novo* single nucleotide variants in BD and SCZ has been investigated in only a handful of studies so far, enrichment in pathways associated with the postsynaptic density has been reported for SCZ, but not BD^14,15^. Identifying disorder-specific variants or quantifying the contribution of variation to specific symptom dimensions remains an open question. For example, previous work by this group has demonstrated that SCZ patients with greater manic symptoms had higher polygenic risk for BD^7^. Here, we utilize the largest collection of genotyped samples of BD and SCZ along with 28 subphenotypes to assess variants and genomic regions that contribute differentially to the disorders and to specific symptoms dimensions or subphenotypes within them.

## Methods

### Sample Description

SCZ samples are those analyzed previously^16^. BD samples are the newest collection from Psychiatric Genomics Consortium Bipolar Disorder Working Group (*Stahl et al. submitted*). To ensure independence of the data sets, individuals were excluded until no individual showed a relatedness (pihat) value greater than 0.2 to any other individual in the collection, while preferentially keeping the case over the control for case-control related pairs. In total 2,181 BD cases, 1,604 SCZ cases and 27,308 controls were removed (most of which were previously known), leaving 20,129 BD cases 33,426 SCZ cases and 54,065 controls for the final metaanalysis.

For analyses directly comparing BD and SCZ, we matched cases from both phenotypes on genotyping platform and ancestry, resulting in 15,270 BD cases versus 23,585 SCZ cases. In other words, we were able to match 76% of BD cases and 71% of SCZ cases.

### Sub-phenotype description

BD sub-phenotypes were collected by each study site using a combination of diagnostic instruments, case records and participant interviews. Ascertainment details for each study site are described in the supplementary data of the PGC Bipolar Working Group paper (*Stahl et al. submitted*). The selection of phenotypes for collection by this group was determined by literature searches in order to determine phenotypes with prior evidence for heritability. It was further refined dependent on the availability of phenotype data across a range of study sites and the consistency by which the phenotypes were defined. Schizophrenia subphenotypes are the same as described previously but in a larger proportion of patients^7^.

### Quality Control, Imputation, Association Analysis and Polygenic Risk Scoring

Quality control and imputation were performed on each of the study cohort datasets (n=81), according to standards established by the Psychiatric Genomics Consortium (PGC). The quality control parameters for retaining SNPs and subjects were: SNP missingness < 0.05 (before sample removal); subject missingness (p < 0.02); autosomal heterozygosity deviation (| F_he_t | < 0.2); SNP missingness < 0.02 (after sample removal); difference in SNP missingness between cases and controls < 0.02; and SNP Hardy-Weinberg equilibrium (p > 10^−6^ in controls or p > 10^−10^ in cases). Genotype imputation was performed using the pre-phasing/imputation stepwise approach implemented in IMPUTE2^17^ / SHAPEIT^18^ (chunk size of 3 Mb and default parameters). The imputation reference set consisted of 2,186 phased haplotypes from the full 1000 Genomes Project dataset (August 2012, 30,069,288 variants, release “v3.macGT1”). After imputation, we used the best guess genotypes, for further robust relatedness testing and population structure analysis. Here we required very high imputation quality (INFO > 0.8) and low missingness (<1%) for further quality control. After linkage disequilibrium (LD) pruning (r^2^ < 0.02) and frequency filtering (MAF > 0.05), there were 14,473 autosomal SNPs in the data set. Relatedness testing was done with PLINK^19^ and pairs of subjects with pihat > 0.2 were identified and one member of each pair removed at random after preferentially retaining cases over controls. Principal component estimation was done with the same collection of autosomal SNPs. We tested the first 20 principal components for phenotype association (using logistic regression with study indicator variables included as covariates) and evaluated their impact on the genomewide test statistics using λ. Thirteen principal components namely 1,2,3,4,5,6,7,8,10,12,15,18,20 were included in all association analyses (λ=1.45). Analytical steps were repeated for SCZ vs BD analysis.

We performed four main association analyses, i.e. (i) GWAS of BD and SCZ as a single combined case phenotype, as well as disorder-specific GWAS using independent control sets in (ii) BD cases vs BD controls and (iii) SCZ cases vs SCZ controls, and (iv) association analysis of SCZ cases vs BD cases.

### Summary-data-based Mendelian Randomization (SMR)^20^

We used SMR as a statistical fine-mapping tool applied to the SCZ vs BD GWAS results to identify loci with strong evidence of causality via gene expression. SMR analysis is limited to significant (FDR < 0.05) cis SNP-expression quantitative trait loci (eQTLs) with MAF > 0.01. eQTLs passing these thresholds were combined with GWAS results in the SMR test, with significance (p_SMR_) reported at a Bonferroni-corrected threshold for each eQTL data set. The eQTL architecture may differ between genes. Through LD, many SNPs can generate significant associations with the same gene, but in some instances multiple SNPs may be independently associated with the expression of a gene. After identification of significant SNP-expression-trait association through the SMR test, a follow-up heterogeneity test aims to prioritize variants by excluding regions for which there is conservative evidence for multiple causal loci (p_HET_ < 0.05). SMR analyses were conducted using eQTL data from whole peripheral blood^21^, dorsolateral prefrontal cortex generated by the CommonMind Consortium^8^, and 11 brain sub-regions from the GTEx consortium^22^.

### Regional joint GWAS

Summary statistic Z-scores were calculated for each marker in each of the four main GWAS results, using the logistic regression coefficient and its standard error. Rare SNPs (MAF < 0.01), and SNPs with a low INFO score (< 0.3) in either dataset were removed. The causal variant relationships between SCZ and BD were investigated using the Bayesian method software pw-gwas (v0.2.1), with quasi-independent regions determined by estimate LD blocks in an analysis of European individuals (n=1,702)^23,24^. Briefly, pw-gwas takes a Bayesian approach to determine the probability of five independent models of association. (1) There is no causal variant in BD or SCZ; (2) a causal variant in BD, but not SCZ (3); a causal variant in SCZ, but not BD; (4) a shared causal variant influencing both BD and SCZ; (5) two causal variants where one influences BD, and one influences SCZ. The posterior probability of each model is calculated using model priors, estimated empirically within pw-gwas. Regions were considered to support a particular model when the posterior probability of the model was greater than 0.5.

### Regional SNP-heritability estimation

We calculated local SNP-heritability independently for SCZ and BD using the Heritability Estimator from Summary Statistics (HESS) software^25^ for each of the independent regions defined above. The sum of these regional estimates is the total SNP-heritability of the trait. To calculate local SNP-heritability HESS requires reference LD matrices representative of the population from which the GWAS samples were drawn. We utilized the 1000 genomes European individuals as the reference panel^26^. Unlike pw-gwas^23^, HESS does not assume that only one causal variant can be present in each region.

## Results

### GWAS

We performed association analysis of BD and SCZ as a combined phenotype, totaling 53,555 cases (20,129 BD, 33,426 SCZ) and 54,065 controls on 15.5 million dosages imputed from 1000 genomes phase 3^26^. Logistic regression was performed controlling for 13 components of ancestry, study sites and genotyping platform. One hundred and fourteen regions contained at least one variant for which the p-value was lower than our genome-wide significance (GWS) threshold of p < 5×10^−8^. Among these 114 loci, 41 had non-overlapping LD regions (r^2^ > 0.6) with the largest and most recently performed single disease GWAS of SCZ^16^ and BD (*Stahl et al. submitted*). Establishing independent controls (see Methods) allowed us to perform disorder-specific GWAS in 20,129 BD cases vs 21,524 BD controls and 33,426 SCZ cases and 32,541 SCZ controls. Using these results, we compared effect sizes of these 114 loci across each disorder independently (Figure 1a) showing that subsets of variants have larger effects in SCZ vs BD or vice versa.

**Figure 1.**
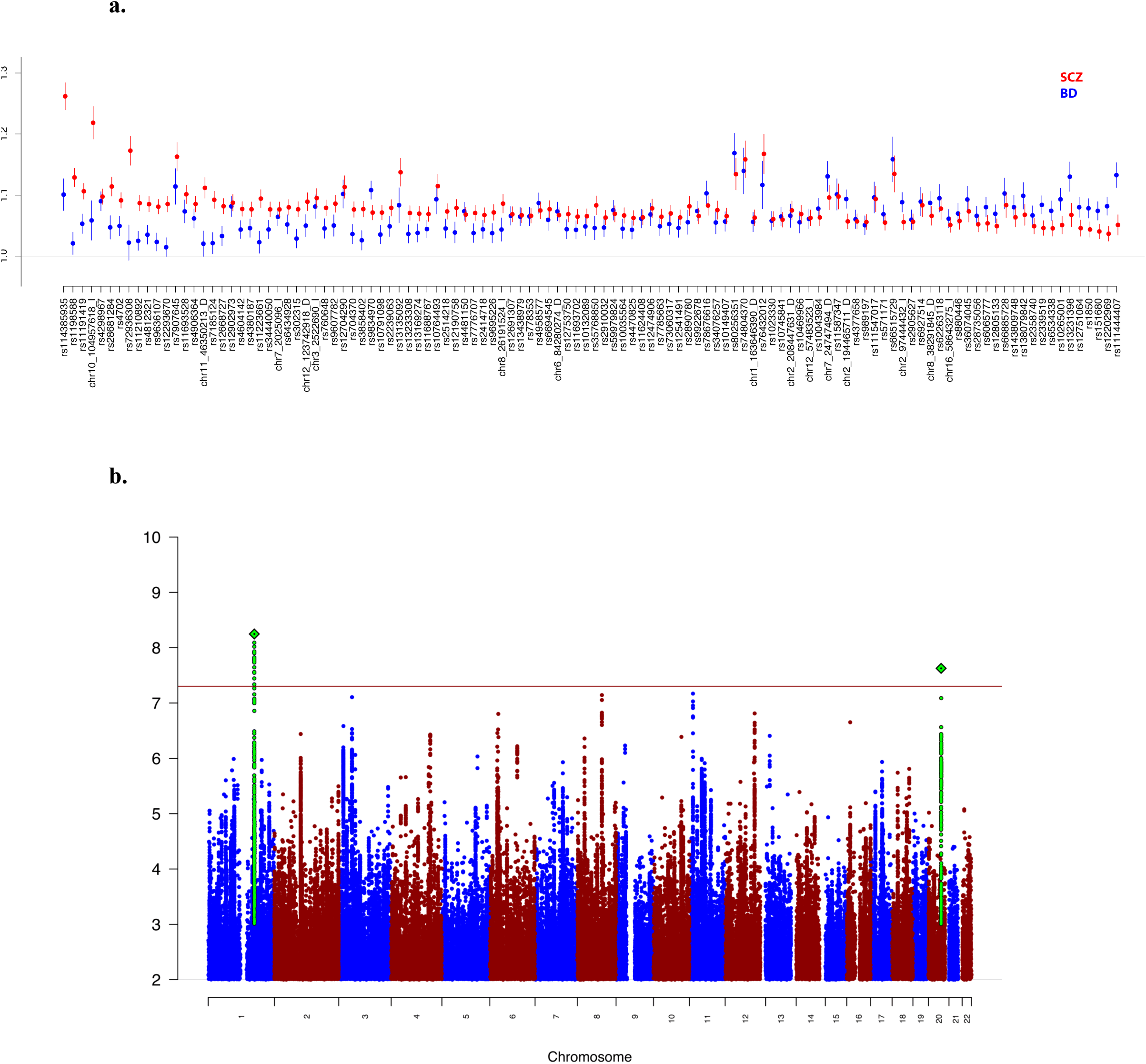
a) Odds ratios (OR) from independent data sets of BD (blue) and SCZ (red) for each of the 114 genome-wide significant variants in the BD and SCZ vs controls GWAS. b) Manhattan plot for SCZ vs BD GWAS.

To identify loci with divergent effects on BD and SCZ, we performed an association analysis on 23,585 SCZ cases and 15,270 BD cases matched for shared ancestry and genotyping platform (see Methods, Figure 1b Supplementary Figures 1-5, Supplementary Table 1). Two genomewide significant loci were identified, the most significant of which was rs56355601 located on chromosome 1 at position 173,811,455 within an intron of *DARS2.* The second most significant locus was a four base indel on chromosome 20 at position 47638976 in an intron of *ARFGEF2.* For both variants, the minor allele frequency was higher in BD cases than SCZ cases and disease-specific GWAS showed opposite directions of effect. We sought to identify additional disease specific loci by incorporating expression information with association results to perform fine-mapping and identify novel variants^27–30^. Here, we applied the summary-data-based Mendelian randomization (SMR) method^20^ (see Methods) utilizing the cis-QTLs derived from peripheral blood^21^, human dorsolateral prefrontal cortex (DLPFC)^31^ from the Common Mind Consortium and 11 brain regions from the GTEx consortium^22^. We identified one SNP-probe combination that surpassed the threshold for genome-wide significance in blood but was also the most significant finding in brain. We found that SNP rs4793172 in gene *DCAKD* is associated with SCZ vs BD analysis (p_GWAS_ = 2.8×10^−6^) and is an eQTL for probe ILMN 1811648 (p_eQTL_ = 2.9×10^−168^), resulting in p_SMR_ = 4.1×10^−6^ in blood (p_eQTL_ = 2.9×10^−25^, p_SMR_ = 2.0×10^−5^ in DLFC, and p_eQTL_ = 4.6×10^−15^, p_SMR_ = 6.0×10^−5^ in GTEx cerebellar hemisphere) (Supplementary Table 2, Supplementary Figure 6) and shows no evidence of heterogeneity (p_HET_ =0.66) which implies only a single causal variant in the region.

### Regional joint association

We expanded our efforts to identify disorder specific genomic regions by jointly analyzing independent GWAS results from BD and SCZ^23^. Among 1,702 regions genome-wide (see Methods), 223 had a posterior probability of greater than 0.5 of having a causal variant in at least one disorder. Of these, 132 best fit the model of a shared causal variant influencing both BD and SCZ, 88 were most likely specific to SCZ, 3 demonstrated evidence of two independent variants (with one impacting each of the two disorders) and zero were BD specific. Of note, the data estimated prior probability of having a BD specific region was 0.1% compared to 15% for SCZ, potentially a result of increased power from the larger SCZ sample size.

The 114 GWS SNPs from the combined BD and SCZ GWAS localized into 99 independent regions, of which 78 (79%) were shared with a posterior probability of greater than 0.5. Sixty regions had at least one GWS SNP in the independent SCZ GWAS, of which 30 (50%) are shared and 8 regions contained a GWS SNP in the independent BD GWAS, of which 6 (75%) are shared using the same definition. For the three regions showing evidence for independent variants, two had highly non-overlapping association signals in the same region stemming from independent variants. The third, on chromosome 19 presented a different scenario where association signals were overlapping (Supplementary Figure 7). The most significant variant in BD was rs111444407 (chr19:19358207, p = 8.67×10^−10^) and for SCZ was rs2315283 (chr19:19480575, p=4.41×10^−7^). After conditioning on the most significant variant in the other disorder, the association signals of the most significant variant in BD and SCZ were largely unchanged (BD rs111444407 =1.3×10^−9^, SCZ rs2315283 p=6.7×10^−5^). We further calculated the probability of each variant in the region being causal for both BD and SCZ^32^ and found no correlation (r= -0.00016). The most significant variants had the highest posterior probability of being causal (SCZ: rs2315283, prob = 0.02, BD: rs111444407, prob = 0.16). Both variants most significantly regulate the expression of *GATAD2A* in brain^31^ but in opposite directions (rs111444407 p_eQTL_ = 6×10^−15^, beta = 0.105; rs2315283 p_eQTL_ = 1.5×10^−28^, beta = -0.11).

### Regional SNP-heritability estimation

Across the genome, regional SNP-heritabilities (h^2^_snp_) were estimated separately for SCZ and BD^25^ and were found to be moderately correlated (r=0.25). We next defined risk regions as those containing the most associated SNP for each GWS locus. In total, there were 101 SCZ risk regions from the 105 autosomal GWS loci reported previously^16^ and 29 BD risk regions from 30 GWS loci reported in a companion paper (*Stahl et al. submitted*). Ten regions were risk regions for both BD and SCZ comprising 33% of BD risk regions and 10% of SCZ risk regions. We further stratified regional h^2^_snp_ by whether a region was a risk region in one disorder, none or both (Figure 2). Since the discovery data for the regions overlapped with the data used for the heritability estimation, we expected within-disorder analyses to show significant results. In risk regions specific to SCZ (n=91) there was a significant increase in regional h^2^_snp_ in SCZ, as expected (p = 1.1×10^−22^), but also in BD (p = 1.2×10^−6^). In risk regions specific to BD (n=19), significantly increased regional h^2^_snp_ was observed in BD, as expected (p = 0.0007), but not in SCZ (p = 0.89). Risk regions shared by both disorders had significantly higher h^2^_snp_ in both disorders, as expected (BD p = 5.3×10^−5^, SCZ p = 0.006), compared to non-risk regions. However, we observed a significant increase in BD h^2^_snp_ in shared risk regions compared to BD risk regions (BD p = 0.003) but not SCZ h^2^_snp_ for shared risk regions compared to SCZ risk regions (p = 0.62). Using a less stringent p-value threshold for defining risk regions (p < 5×10^−6^), thereby substantially increasing the number of regions, resulted in similar results (Supplementary Figure 8). Seven regions contributed to substantially higher h^2^_snp_ in SCZ compared to BD but no region showed the inverse pattern. Of these regions, all but one was in the major histocompatibility region (MHC), the sole novel region was chr10:104380410-106695047 with regional h^2^_snp_= 0.0019 in SCZ and h^2^_snp_=0.00063 in BD.

**Figure 2.**
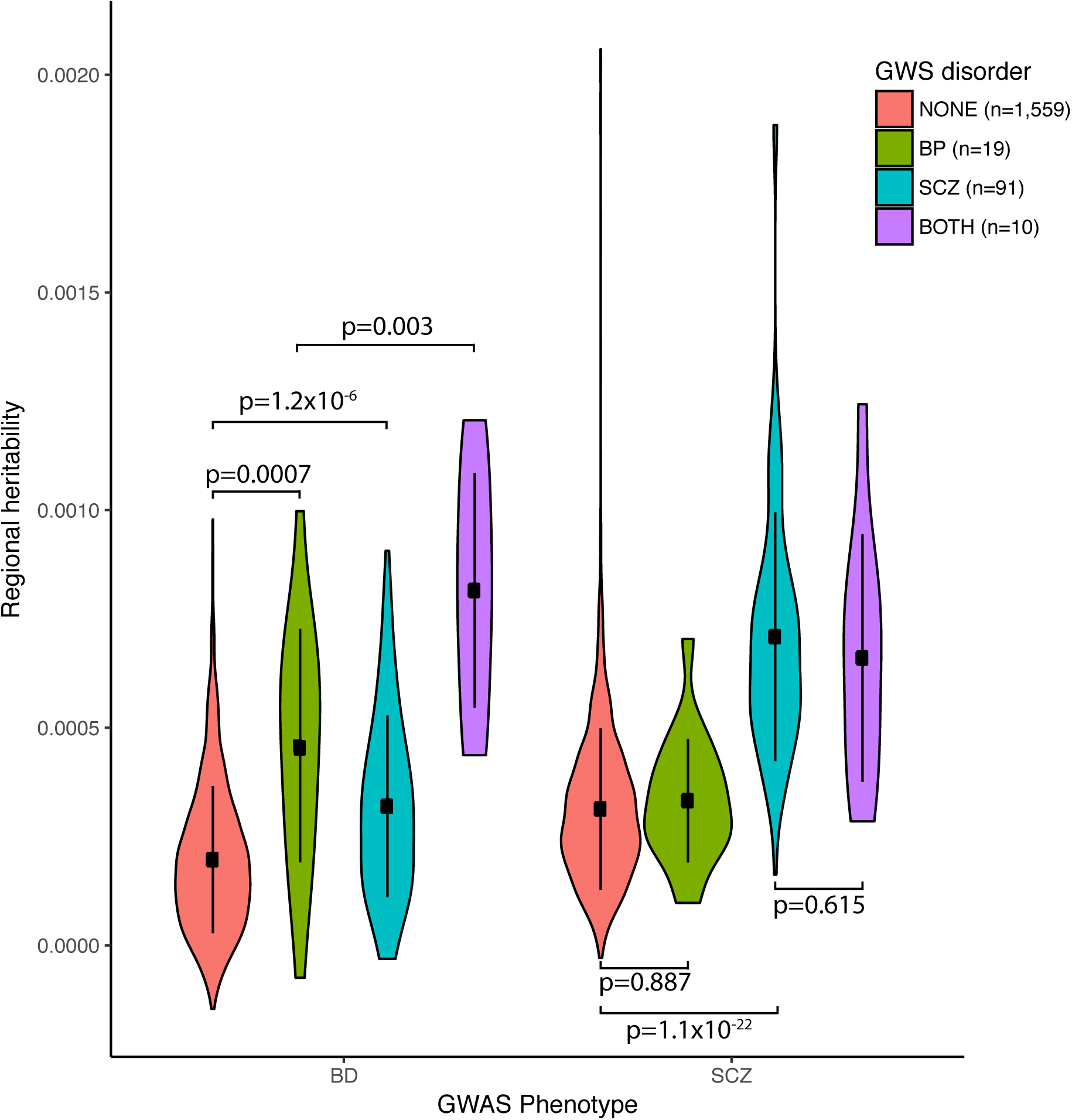
Regional SNP-heritability estimates for SCZ and BD stratified by whether the region contains the most significant variant in a genome-wide significant locus in BD, SCZ, neither or both.

### Polygenic dissection of subphenotypes

subphenotypes were collected for a subset of patients in both BD and SCZ (see Methods). For SCZ, we had clinical quantitative measurements of manic, depressive, positive and negative symptoms generated from factor analysis of multiple instruments as described previously^7^ but in larger sample sizes (n=6908, 6907, 8259, 8355 respectively). For BD, 24 subphenotypes were collected among nearly 13,000 cases in distinct categories including comorbidities, clinical information such as rapid cycling and psychotic features as well as additional disease course data such as age of onset and number of hospitalizations. For each BD and SCZ patient, we calculated a polygenic risk score (PRS) using all SNPs, from each of the four main GWAS analyses (BD+SCZ, BD, SCZ and SCZvsBD). We then used regression analysis including principal components and site to assess the relationship between each subphenotype and the 4 PRS. We applied a significance cutoff of p < 0.0004 based on Bonferroni correction for 112 tests. In total, we identified 6 significant results after correction (Figure 3, Table 1). For BD PRS we see a significant positive correlation between PRS and manic symptoms in SCZ cases as seen previously^7^ (p=2×10^−5^, t=4.26) and psychotic features in BD patients (p=5.3×10^−5^, t=4.04). For SCZ PRS, we see a significant increase in PRS for BD cases with versus without psychotic features (p=1.2×10^−10^, t=6.45) and negative symptoms in SCZ patients (p=3.60×10^−6^, t=4.64). As with the SCZ PRS, BD+SCZ PRS is also significantly associated with psychotic features in BD (p=7.9×10^−13^, t=7.17) and negative symptoms in SCZ (p=1.5×10^−5^, t=4.33). While not surpassing conservative correction, the next two most significant results are both indicative of a more severe course in BD: increased BD+SCZ PRS with increased numbers of hospitalizations in BD cases (p=4.2×10^−4^, t=3.53) and increased SCZ PRS with earlier onset of BD (p=7.9×10^−4^, t=-3.36). We assessed the role of BD subtype on correlation between SCZ PRS and psychotic features and identified significant correlation when restricted to only BD type I cases (BDI: 3,763 with psychosis, 2,629 without, p=1.55×10^−5^, Supplementary Table 3).

**Figure 3.**
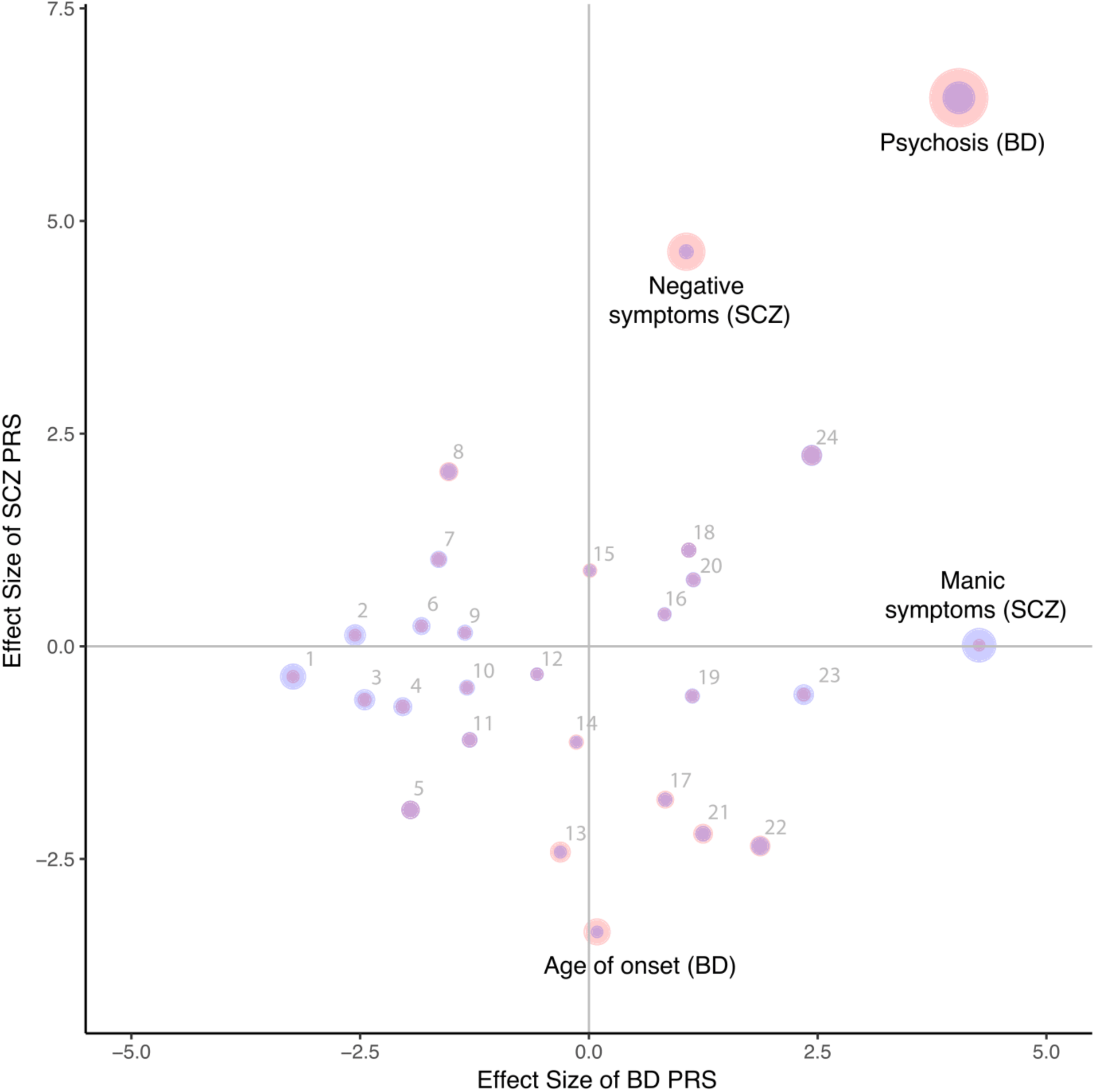
Effect size (calculated by dividing regression estimate by standard error) from regression analysis including ancestry covariates for each subphenotype and PRS for BD (x-axis) and SCZ (y-axis). Point size represents –log10(p-value) with SCZ (red) and BD (blue). Numbered subphenotypes are 1) comorbid migraine, 2) panic attacks 3) suicide attempt 4) mixed states 5) rapid cycling 6) comorbid eating disorder 7) comorbid OCD 8) year of birth 9) suicide ideation 10) panic disorder 11) number of suicide attempts 12) depressive symptoms (SCZ) 13) episodes depressive 14) episodes total 15) positive symptoms (SCZ) 16) irritable mania 17) age of onset depression 18) family history 19) episodes mixed mania 20) unipolar mania 21) alcohol substance dependence 22) age of onset mania 23) age at interview 24) number of hospitalizations. All subphenotypes are in BD except those labeled (SCZ).

**Table 1.** Polygenic scoring results of all four GWAS phenotypes (BD+SCZ vs controls, BD vs controls, SCZ vs controls and SCZ vs BD) and 24 subphenotypes from BD and 4 subphenotypes from SCZ, rows without case/control counts are quantitative measures. Significance and effects are from regression analysis of subphenotype on PRS including ancestry and site as covariates. Effect is the regression estimate divided by the standard error.

|  |  |  |  |  | P-value |  |  |  | Effect |  |  |  |
| --- | --- | --- | --- | --- | --- | --- | --- | --- | --- | --- | --- | --- |
|  | Subphenotype | N | Cases | Controls | BP+SCZ | BP | SCZ | SCZvsBD | BP+SCZ | BP | SCZ | SCZvsBD |
| BD | psychosis | 8131 | 4632 | 3499 | 7.9E-13 | 5.3E-05 | 1.2E-10 | 5.8E-01 | 7.17 | 4.04 | 6.45 | 0.55 |
|  | suicide ideation | 5399 | 3801 | 1598 | 7.8E-01 | 1.8E-01 | 8.7E-01 | 1.7E-01 | -0.28 | -1.35 | 0.16 | 1.37 |
|  | family history | 4971 | 2730 | 2241 | 6.1E-02 | 2.8E-01 | 2.6E-01 | 6.9E-01 | 1.87 | 1.09 | 1.13 | -0.39 |
|  | irritable mania | 4230 | 2401 | 1829 | 3.8E-01 | 4.1E-01 | 7.1E-01 | 1.0E-01 | 0.88 | 0.83 | 0.38 | -1.63 |
|  | rapid cycling | 5214 | 1744 | 3470 | 7.9E-03 | 5.1E-02 | 5.5E-02 | 3.1E-01 | -2.66 | -1.95 | -1.92 | 1.01 |
|  | alcohol substance dependence | 5440 | 1494 | 3946 | 4.5E-01 | 2.1E-01 | 2.8E-02 | 1.7E-01 | -0.75 | 1.25 | -2.20 | -1.36 |
|  | panic disorder | 4647 | 863 | 3784 | 2.8E-01 | 1.8E-01 | 6.3E-01 | 4.0E-01 | -1.07 | -1.33 | -0.49 | 0.83 |
|  | panic attacks | 3976 | 851 | 3125 | 1.3E-01 | 1.1E-02 | 9.0E-01 | 4.7E-02 | -1.50 | -2.56 | 0.13 | 1.98 |
|  | mixed states | 4044 | 826 | 3218 | 1.0E-01 | 4.2E-02 | 4.8E-01 | 6.0E-02 | -1.64 | -2.03 | -0.71 | 1.88 |
|  | unipolar mania | 4863 | 461 | 4402 | 2.4E-02 | 2.5E-01 | 4.3E-01 | 6.1E-01 | 2.26 | 1.14 | 0.78 | 0.51 |
|  | comorbid migraine | 2652 | 410 | 2242 | 1.3E-02 | 1.2E-03 | 7.2E-01 | 4.4E-01 | -2.48 | -3.23 | -0.36 | 0.77 |
|  | comorbid OCD | 4215 | 386 | 3829 | 9.7E-01 | 1.0E-01 | 3.1E-01 | 1.9E-01 | -0.04 | -1.64 | 1.02 | 1.30 |
|  | comorbid eating disorder | 3839 | 331 | 3508 | 2.1E-01 | 6.7E-02 | 8.1E-01 | 6.3E-01 | -1.25 | -1.83 | 0.24 | 0.48 |
|  | age of onset | 8610 |  |  | 6.2E-03 | 9.3E-01 | 7.9E-04 | 6.2E-01 | -2.74 | 0.09 | -3.36 | -0.50 |
|  | age at interview | 8062 |  |  | 5.9E-01 | 1.9E-02 | 5.7E-01 | 4.4E-01 | 0.54 | 2.35 | -0.57 | -0.78 |
|  | episodes mixed mania | 6587 |  |  | 6.3E-01 | 2.6E-01 | 5.6E-01 | 3.2E-01 | -0.48 | 1.13 | -0.58 | -1.00 |
|  | suicide attempt | 6308 |  |  | 1.2E-01 | 1.4E-02 | 5.3E-01 | 2.8E-01 | -1.54 | -2.45 | -0.63 | 1.09 |
|  | episodes depressive | 6252 |  |  | 7.4E-03 | 7.6E-01 | 1.6E-02 | 9.6E-01 | -2.68 | -0.31 | -2.42 | -0.05 |
|  | episodes total | 5958 |  |  | 1.3E-01 | 8.9E-01 | 2.6E-01 | 3.9E-01 | -1.51 | -0.14 | -1.13 | -0.87 |
|  | year of birth | 5317 |  |  | 1.7E-01 | 1.3E-01 | 4.0E-02 | 3.6E-02 | 1.39 | -1.53 | 2.05 | 2.10 |
|  | number of suicide attempts | 5015 |  |  | 6.2E-02 | 1.9E-01 | 2.7E-01 | 4.9E-01 | -1.87 | -1.30 | -1.10 | -0.69 |
|  | number of hospitalizations | 3944 |  |  | 4.2E-04 | 1.5E-02 | 2.5E-02 | 7.4E-01 | 3.53 | 2.43 | 2.25 | -0.33 |
|  | age of onset depression | 3467 |  |  | 2.3E-01 | 4.0E-01 | 7.2E-02 | 2.2E-01 | -1.19 | 0.83 | -1.80 | 1.24 |
|  | age of onset mania | 3395 |  |  | 2.5E-01 | 6.1E-02 | 1.9E-02 | 2.2E-01 | -1.14 | 1.87 | -2.35 | -1.23 |
| SCZ | Manic | 6908 |  |  | 2.4E-02 | 2.0E-05 | 9.9E-01 | 3.5E-02 | 2.26 | 4.26 | 0.01 | -2.10 |
|  | Depressive | 6907 |  |  | 9.0E-01 | 5.7E-01 | 7.4E-01 | 1.8E-01 | 0.13 | -0.57 | -0.33 | -1.36 |
|  | Negative | 8355 |  |  | 1.5E-05 | 2.9E-01 | 3.6E-06 | 2.1E-02 | 4.33 | 1.06 | 4.64 | 2.31 |
|  | Positive | 8259 |  |  | 4.1E-01 | 9.9E-01 | 3.7E-01 | 5.1E-01 | 0.82 | 0.01 | 0.89 | 0.65 |

For all 8 quantitative subphenotypes and 9 binary subphenotypes having at least 1,000 cases, we performed a GWAS within cases to calculate heritability and genetic correlation with BD and SCZ. Only two subphenotypes had significant h^2^_snp_ estimates using LD-score regression^33^, psychotic features in BD (h^2^_snp_=0.15, SE=0.06) and suicide attempt (h^2^_snp_=0.25, SE=0.1). Only psychotic features demonstrated significant genetic correlation with SCZ (r_g_=0.34, SE=0.13, p=0.009). While the genetic correlation demonstrates a genome-wide relationship between common variants contributing to SCZ and those contributing to psychotic features in BD cases, we sought to assess whether this could be demonstrated among the most significantly associated SCZ loci. Of the 105 autosomal genome-wide significant SCZ loci previously published^16^, 60 out of 100 variants in our dataset after QC demonstrated the same direction of effect for psychotic features in BD (p=0.028, one-sided binomial-test).

## Discussion

Here we present a genetic dissection of bipolar disorder and schizophrenia from over 100,000 genotyped subjects. As previously shown^34^, we found an extensive degree of genetic sharing between these two disorders. We identified 114 genome-wide significant loci contributing to both disorders of which 37 are novel to this analysis. Despite the high degree of sharing, we identified several loci that significantly differentiated between the two disorders, having opposite directions of effect, and polygenic components that significantly correlated from one disorder to symptoms of the other.

Two GWS loci were identified from the case only SCZ versus BD analysis providing opportunities to inform the underlying biological distinctions between BD and SCZ. The most significant locus is in *DARS2* (coding for the mitochondrial Aspartate-tRNA ligase) which is highly expressed in the brain and significantly regulated by the most significant SNP rs56355601 (p_eQTL_=2.5×10^−11^). Homozygous mutations in *DARS2* are responsible for leukoencephalopathy with brainstem and spinal cord involvement and lactate elevation (LBSL), which was characterized by neurological symptoms such as psychomotor developmental delay, cerebellar ataxia and delayed mental development^35^. Interestingly, based on methylation analysis from the prefrontal cortex of stress models (rats and monkeys) and from peripheral samples (in monkeys and human newborns), *DARS2*, among others, has been suggested as a potential molecular marker of early-life stress and vulnerability to psychiatric disorders^36^. The second most significant locus maps to *ARFGEF2*, which codes for ADP Ribosylation Factor Guanine Nucleotide Exchange Factor 2 (also known as BIG2), a protein involved in vesicular trafficking from the trans-Golgi network. Mutations in *ARFGEF2* have been shown to underlie an autosomal recessive condition characterized by microcephaly and periventricular heterotopia, a disorder caused by abnormal neural proliferation and migration^37^. Although not genome-wide significant, the third most significant locus implicates *ARNTL* (Aryl Hydrocarbon Receptor Nuclear Translocator Like), which is a core component of the circadian clock. *ARNTL* has been previously hypothesized for relevance in bipolar disorder,^38^ although human genetic evidence is limited^39^. Incorporating transcriptional data identified a third genome-wide significant finding in *DCAKD.* The gene codes for Dephospho-CoA Kinase Domain Containing, a member of the human postsynaptic density proteome from human neocortex^40^. In the mouse cortical synaptoproteome *DCAKD* has been found to be among the proteins with the highest changes between juvenile postnatal days and adult stage, which suggests a putative role in brain development^41,42^.

We further assessed the contribution of regions of the genome to each disorder through joint regional association and regional heritability estimation. These results point to two additional loci that may contribute differentially to liability to BD and SCZ. The region on chr19 shows overlapping association peaks that are driven by independent causal variants for each disorder. Both variants significantly regulate the same gene *GATAD2A* but in opposite directions. *GATAD2A* is a transcriptional repressor, which is targeted by *MBD2* and is involved in methylation-dependent gene silencing. The protein is part of the large NuRD (nucleosome remodeling and deacetylase) complex, for which also HDAC1/2 are essential components. NurD complex proteins have been associated to autism^43^. Their members, including *GATAD2A*, display preferential expression in fetal brain development^43^ and in recent work has been implicated in SCZ through open chromatin^44^. Further, p66α (mouse *GATAD2A*) was recently shown to participate in memory preservation through long-lasting histone modification in hippocampal memory-activated neurons^45^. The region on chromosome 10 appears to be shared across both disorders; however, there are additional independent contributing variants to SCZ and not BD, indicating another region of interest, although biological interpretation remains unknown.

More broadly, SNP-heritability appears to be consistently shared across regions and chromosomes between these two disorders. Regions with GWS loci often explain higher proportions of heritability as expected. When looking at the effect on heritability of the presence of a GWS locus in the other disorder, we identified a significant increase in BD heritability for regions containing a GWS locus for SCZ but no significant increase in SCZ heritability in regions having a BD one. This result suggests a directionality to the genetic sharing of these disorders with a larger proportion of BD loci being specific to BD. However, we cannot exclude that the asymmetry of results may reflect less power of discovery for BD than SCZ. The degree to which power and subphenotypes contribute to this result requires further examination.

We have now identified multiple genomic signatures that correlate between one disorder and a clinical symptom in the other disorder, demonstrating that there are genetic components underlying particular symptom dimensions within these disorders. As previously shown, we find a significant positive correlation between PRS of BD and manic symptoms in SCZ. We also demonstrate that BD cases with psychotic features carry a significantly higher SCZ PRS than BD cases without psychotic features and this result is not driven by schizoaffective BD subtype. Further, we show evidence that increased PRS is associated with more severe illness. This is true for BD with psychotic features having increased SCZ PRS, earlier onset BD having higher SCZ PRS and cases with higher BD+SCZ PRS having a larger number of hospitalizations. We demonstrated that psychotic features within BD is an independently heritable trait and that GWS loci for SCZ have a consistent direction of effect in psychotic features in BD, demonstrating the potential to study psychosis more directly to identify variants contributing to that symptom dimension. All in all, this work illustrates the utility of genetic data to dissect symptom heterogeneity among correlated disorders and suggests that further work could potentially aid in defining subgroups of patients for more personalized treatment.

## References

1. Whiteford, H. A. et al. Global burden of disease attributable to mental and substance use disorders: findings from the Global Burden of Disease Study 2010. The Lancet 382, 1575–1586 (2013).

2. Nöthen, M. M., Nieratschker, V., Cichon, S. & Rietschel, M. New findings in the genetics of major psychoses. Dialogues Clin. Neurosci. 12, 85–93 (2010).

3. Lichtenstein, P. et al. Common genetic determinants of schizophrenia and bipolar disorder in Swedish families: a population-based study. The Lancet 373, 234–239 (2009).

4. Diagnostic and Statistical Manual of Mental Disorders | DSM Library. Available at: http://dsm.psychiatryonline.org/doi/book/10.n76/appi.books.9780890425596. (Accessed: 14th March 2017)

5. WHO | International Classification of Diseases. WHO Available at: http://www.who.int/classifications/icd/en/. (Accessed: 14th March 2017)

6. Bortolato, B., Miskowiak, K. W., Köhler, C. A., Vieta, E. & Carvalho, A. F. Cognitive dysfunction in bipolar disorder and schizophrenia: a systematic review of meta-analyses. Neuropsychiatr. Dis. Treat. 11, 3111–3125 (2015).

7. Ruderfer, D. M. et al. Polygenic dissection of diagnosis and clinical dimensions of bipolar disorder and schizophrenia. Mol. Psychiatry 19, 1017–1024 (2014).

8. Kirov, G. et al. De novo CNV analysis implicates specific abnormalities of postsynaptic signalling complexes in the pathogenesis of schizophrenia. Mol. Psychiatry 17, 142–153 (2012).

9. Szatkiewicz, J. P. et al. Copy number variation in schizophrenia in Sweden. Mol. Psychiatry 19, 762–773 (2014).

10. CNV and Schizophrenia Working Groups of the Psychiatric Genomics Consortium. Contribution of copy number variants to schizophrenia from a genome-wide study of 41,321 subjects. Nat. Genet. 49, 27–35 (2017).

11. Stone, J. L. et al. Rare chromosomal deletions and duplications increase risk of schizophrenia. Nature 455, 237–241 (2008).

12. Gulsuner, S. & McClellan, J. M. Copy Number Variation in Schizophrenia. Neuropsychopharmacology 40, 252–254 (2015).

13. Green, E. K. et al. Copy number variation in bipolar disorder. Mol. Psychiatry 21, 89–93 (2016).

14. Kataoka, M. et al. Exome sequencing for bipolar disorder points to roles of de novo loss-of-function and protein-altering mutations. Mol. Psychiatry 21, 885–893 (2016).

15. Fromer, M. et al. De novo mutations in schizophrenia implicate synaptic networks. Nature 506, 179–184 (2014).

16. Schizophrenia Working Group of the Psychiatric Genomics Consortium. Biological insights from 108 schizophrenia-associated genetic loci. Nature 511, 421–427 (2014).

17. Howie, B., Marchini, J. & Stephens, M. Genotype Imputation with Thousands of Genomes. G3 GenesGenomesGenetics 1, 457–470 (2011).

18. Delaneau, O., Zagury, J.-F. & Marchini, J. Improved whole-chromosome phasing for disease and population genetic studies. Nat. Methods 10, 5–6 (2013).

19. Purcell, S. et al. PLINK: A Tool Set for Whole-Genome Association and Population-Based Linkage Analyses. Am. J. Hum. Genet. 81, 559–575 (2007).

20. Zhu, Z. et al. Integration of summary data from GWAS and eQTL studies predicts complex trait gene targets. Nat. Genet. 48, 481–487 (2016).

21. Westra, H.-J. et al. Systematic identification of trans eQTLs as putative drivers of known disease associations. Nat. Genet. 45, 1238–1243 (2013).

22. Consortium, T. Gte. The Genotype-Tissue Expression (GTEx) pilot analysis: Multitissue gene regulation in humans. Science 348, 648–660 (2015).

23. Pickrell, J. K. et al. Detection and interpretation of shared genetic influences on 42 human traits. Nat. Genet. 48, 709–717 (2016).

24. Berisa, T. & Pickrell, J. K. Approximately independent linkage disequilibrium blocks in human populations. Bioinformatics 32, 283–285 (2015).

25. Shi, H., Kichaev, G. & Pasaniuc, B. the Genetic Architecture of 30 Complex Traits from Summary Association Data. Am. J. Hum. Genet. 99, 139–153 (2016).

26. The 1000 Genomes Project Consortium. A global reference for human genetic variation. Nature 526, 68–74 (2015).

27. Giambartolomei, C. et al. Bayesian Test for Colocalisation between Pairs of Genetic Association Studies Using Summary Statistics. PLOS Genet. 10, e1004383 (2014).

28. He, X. et al. Sherlock: Detecting Gene-Disease Associations by Matching Patterns of Expression QTL and GWAS. Am. J. Hum. Genet. 92, 667–680 (2013).

29. Gamazon, E. R. et al. A gene-based association method for mapping traits using reference transcriptome data. Nat. Genet. 47, 1091–1098 (2015).

30. Gusev, A. et al. Integrative approaches for large-scale transcriptome-wide association studies. Nat. Genet. 48, 245–252 (2016).

31. Fromer, M. et al. Gene expression elucidates functional impact of polygenic risk for schizophrenia. Nat. Neurosci. 19, 1442–1453 (2016).

32. Benner, C. et al. FINEMAP: efficient variable selection using summary data from genome-wide association studies. Bioinformatics 32, 1493–1501 (2016).

33. Bulik-Sullivan, B. K. et al. LD Score regression distinguishes confounding from polygenicity in genome-wide association studies. Nat. Genet. 47, 291–295 (2015).

34. Cross-Disorder Group of the Psychiatric Genomics Consortium. Genetic relationship between five psychiatric disorders estimated from genome-wide SNPs. Nat. Genet. 45, 984–994 (2013).

35. Yamashita, S. et al. Neuropathology of leukoencephalopathy with brainstem and spinal cord involvement and high lactate caused by a homozygous mutation of DARS2. Brain Dev. 35, 312–316 (2013).

36. Luoni, A. et al. Ankyrin-3 as a molecular marker of early-life stress and vulnerability to psychiatric disorders. Transl. Psychiatry 6, e943 (2016).

37. Sheen, V. L. et al. Mutations in ARFGEF2 implicate vesicle trafficking in neural progenitor proliferation and migration in the human cerebral cortex. Nat. Genet. 36, 69–76 (2004).

38. Yang, S., Van Dongen, H. P. A., Wang, K., Berrettini, W. & Bućan, M. of circadian function in fibroblasts of patients with bipolar disorder. Mol. Psychiatry 14, 143–155 (2008).

39. Byrne, E. M. et al. Testing the role of circadian genes in conferring risk for psychiatric disorders. Am. J. Med. Genet. Part B Neuropsychiatr. Genet. Off. Publ. Int. Soc. Psychiatr. Genet. 0, 254–260 (2014).

40. Bayés, À. et al. Characterization of the proteome, diseases and evolution of the human postsynaptic density. Nat. Neurosci. 14, 19–21 (2011).

41. Moczulska, K. E. et al. Deep and Precise Quantification of the Mouse Synaptosomal Proteome Reveals Substantial Remodeling during Postnatal Maturation. J. Proteome Res. 13, 4310–4324 (2014).

42. Gonzalez-Lozano, M. A. et al. Dynamics of the mouse brain cortical synaptic proteome during postnatal brain development. Sci. Rep. 6, (2016).

43. Li, J. et al. Identification of Human Neuronal Protein Complexes Reveals Biochemical Activities and Convergent Mechanisms of Action in Autism Spectrum Disorders. Cell Syst. 1, 361–374 (2015).

44. Fullard, J. F. et al. Open chromatin profiling of human postmortem brain infers functional roles for non-coding schizophrenia loci. Hum. Mol. Genet. doi:10.1093/hmg/dd×103

45. Ding, X. et al. Activity-induced histone modifications govern Neurexin-1 mRNA splicing and memory preservation. Nat. Neurosci. advance online publication, (2017).

